# Assessing Bias in Gene Expression Omnibus (GEO) Datasets

**DOI:** 10.1101/2024.11.29.622327

**Authors:** Mahnoor N. Gondal

**Author notes:** Correspondence: Mahnoor N. Gondal.

## Abstract

The Gene Expression Omnibus (GEO) is a widely used repository for gene expression data, but its datasets may be subject to biases in terms of diversity, equity, and inclusion. This study aims to assess DEI-related biases in GEO datasets specifically focusing on gender and ethnicity representation. We curated a subset of GEO datasets by applying filters for organism type, sample size, and DEI-related metadata (gender and ethnicity). Following rigorous data extraction and cleaning, we analyzed 211 datasets, evaluating gender balance and ethnicity/race distribution using quantitative metrics. For gender representation, we calculated a gender ratio for each dataset, with a ratio closer to 1 indicating balanced representation. Ethnicity/race representation was assessed using Chi-square goodness-of-fit tests to identify disparities in ethnic distribution. We ranked datasets based on these DEI criteria to identify those with the highest and lowest bias. Our findings revealed that while many datasets displayed balanced gender ratios, significant bias in ethnicity representation was observed, with a predominance of White and African American participants. Additionally, we observed some variation in DEI metrics for datasets published before and after 2015, suggesting a shift in gender recruitment practices. These results highlight the need for more inclusive data collection and reporting in genomic research. We emphasize the importance of incorporating DEI criteria into data curation and encourage future efforts to standardize DEI data reporting across repositories to mitigate bias and improve the generalizability of genomic research.

## Background

The Gene Expression Omnibus (GEO) serves as a vital repository for genomic data, facilitating groundbreaking research across various fields of biology and medicine ^1–5^. However, the representativeness of these datasets in terms of diversity, equity, and inclusion (DEI) remains a critical yet understudied aspect ^6,7^. This study aims to address this gap by systematically assessing DEI-related biases in GEO datasets, with a specific focus on gender and ethnicity representation. By examining these crucial factors, we seek to shed light on potential disparities in genomic data collection and highlight areas for improvement in ensuring more inclusive and generalizable research outcomes.

The importance of diverse and representative genomic data cannot be overstated ^8–10^. Historically, genomic studies have been predominantly conducted on populations of European descent, leading to potential biases in our understanding of genetic variations and their implications for health and disease. This lack of diversity can result in limited applicability of research findings to underrepresented populations and may perpetuate health disparities. As the scientific community increasingly recognizes the need for more inclusive research practices, it becomes crucial to evaluate the current state of diversity in widely used genomic data repositories such as GEO ^11–15^.

Our analysis encompasses a comprehensive evaluation of 211 datasets, employing quantitative metrics to assess gender balance and ethnicity/race distribution. We utilize gender ratios and Chi-square goodness-of-fit tests to quantify representativeness and potential biases in these datasets. Additionally, we examine temporal trends by comparing datasets published before and after 2015 to identify any shifts in data collection practices. By providing insights into the current state of DEI in genomic data repositories, this study aims to contribute to ongoing efforts to improve the inclusivity and generalizability of genomic research.

## Methods

### 1 Data collection

#### Extraction and filtering of GEO data

To extract datasets relevant to our task, we first filtered our search with three initial criteria: (i) type of experiment, (ii) organism, and (iii) sample size. The filters were used directly in the GEO advanced search option. The result from this filtering step generated ∼2563 series. To further make our search more stringent for the US population, we added the USA filter which further reduced our result to 1433 series. While looking at the generated results, we noticed that almost half of the datasets did not contain the DEI criteria under investigation (mainly gender and ethnicity/race). To improve analysis time and efficiency we added a final filter for race and gender for our GEO search. The resultant search showed 582 series. A detail of the steps for data collection is outlined in **Figure 1A**. GEOquery package in R was used for extracting metadata for each data series.

**Figure 1.**
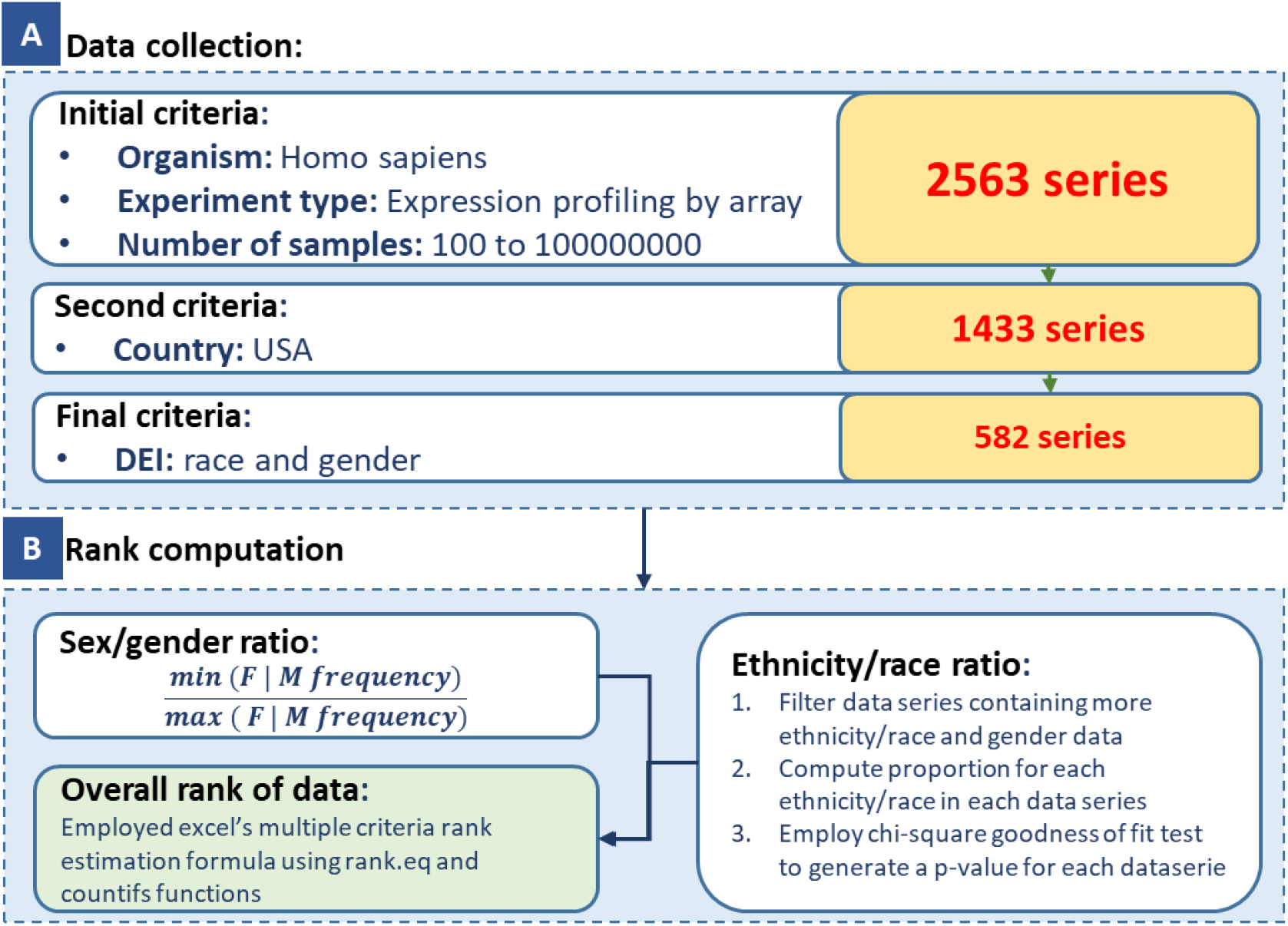
Steps of the analysis

#### Challenges in data collection

Important to note that the metadata in GEO, while comprehensive, lacks uniform structure. When collecting diversity, equity, and inclusion (DEI) information, we observed inconsistencies in reporting biological sex and gender identity. Some datasets used “male” and “female,” while others employed “M” and “F” conventions. Moreover, the terms “sex” and “gender” were often used interchangeably, despite their distinct meanings in DEI contexts. Similarly, we found inconsistencies in reporting ancestry, ethnicity, and race. Some datasets used “ethnicity” labels, while others employed “race.” Abbreviations like “AA” for African Americans and “CAU” for Caucasians were common, often without clarification of their full forms, which required referencing original manuscripts. These inconsistencies highlight the need for standardized reporting practices in GEO metadata. For this study, our primary goal was to evaluate the distribution of male and female subjects in GEO datasets, acknowledging the limitations of the current metadata structure. Additional challenges included poor formatting in some metadata and file size limitations for local processing. Due to these issues, we manually verified each metadata file after downloading to ensure accurate column extraction and data integrity. This labor-intensive process allowed us to analyze 211 out of the 582 filtered data series. Further filtering for specific analysis requirements reduced this number further, as detailed in subsequent sections. These observations underscore the importance of clear, consistent, and comprehensive metadata reporting in genomic databases to facilitate more accurate and inclusive analyses.

### 2 Scoring and rank computation

#### Gender ratio

Towards computing a gender ratio, we employed the following formula, in which out of the total male and female frequency calculated from each series, we used the smaller of the two numbers in the numerator and the larger of the two numbers in the denominator. This approach was undertaken for each data series. This showed that a ratio close to one would be a good ratio (suggesting a balanced gender) whereas a ratio close to zero would be a bad ratio (indicating an unbalanced gender) the total ratio cannot be more than one which came into use with overall rank estimation. An overview of the scoring method can be found in **Figure 1B**. Here, we first calculated the rank of samples based on the gender ratio only and then we incorporated additional criteria and added complexity for computing rank.

#### Ethnicity/race ratio

For calculating the ethnicity/race ratio, we carried out the following steps:

(1) We first filtered those data series with both ethnicity and gender mentioned in their metadata so that we have more of these data for overall rank calculation.
(2) We then calculated the proportion of each ethnicity/race for each data series by dividing each ethnicity/race frequency from the total number. This result gave a reasonable estimate of how much each ethnicity/race is contributing to the overall dataset. While carrying this out, we made sure we were not over-estimating by incorporating ethnicities not mentioned in the respective dataset. Only ethnicities mentioned in the metadata for that series were employed.
(3) We then employed the Chi-square goodness of fit test to estimate a p-value for each data series. This p-value indicates how much is the expected proportion different from the observed proportion of races. A larger p-value indicated more balanced data for ethnicity/race whereas a smaller p-value indicates a more biased dataset. Since failure to reject the null hypothesis concludes that the data does follow a distribution with certain proportions and there is no evidence to suggest that there is a significant difference between observed and expected frequency.

#### Overall rank computation

To estimate the overall rank for each data series we used our prior knowledge:

- For the gender ratio, closer to 1 indicates unbiased
- For the ethnicity/race ratio, a larger p-value indicates an unbiased

Given the way we have structured our results, we know that the gender ratio cannot be larger than 1, and the largest p-value would be the most unbiased for ethnicity/race. Therefore, we used Excel’s internal rank computation formula we first ranked the data series based on the gender ratio and built on that with the ethnicity/race ratio.

## Results

### 1 Investigating bias in Gene Expression Omnibus datasets using DEI criteria

#### Data curated for DEI criteria investigation

To investigate whether Gene expression Omnibus (GEO) datasets are biased relative to the US population using DEI criteria we first added three different filters (i) expression profiling by array, (ii) homo sapiens, and (iii) at least 100 samples. This resulted in around 2563 series. Additional filters for USA and DEI criteria (gender and ethnicity/race) were retrieved around 580 series. Due to poor structure of metadata, downloading issues, missing values, variability in labeling, etc, out of 580 series we manually extracted relevant data for 211 of them - making sure we got the right frequencies.

#### Ranking GEO datasets based on gender

To compute a rank for each dataset based on gender bias, we first calculated a ratio for estimating the proportion of gender. A value close to one indicates a good or balanced dataset whereas a value near zero indicates a bad or unbalanced dataset. Twenty datasets had a ratio of 0 which indicated that they had only one of the genders mentioned. This might be because they focus on a gender-based study such as breast cancer is more prevalent in females whereas male subjects are more prone to prostate cancer. We removed these data series from our results. Some datasets were also removed because they did not contain gender information (they contained race/ethnicity information). The top and bottom five datasets are highlighted in **Figure 2A**. The result shows that there are 116 datasets from 163 which are equal to and above 0.5 indicating a good balance of gender, whereas 43 were below 0.5 showing gender inequality.

**Figure 2.**
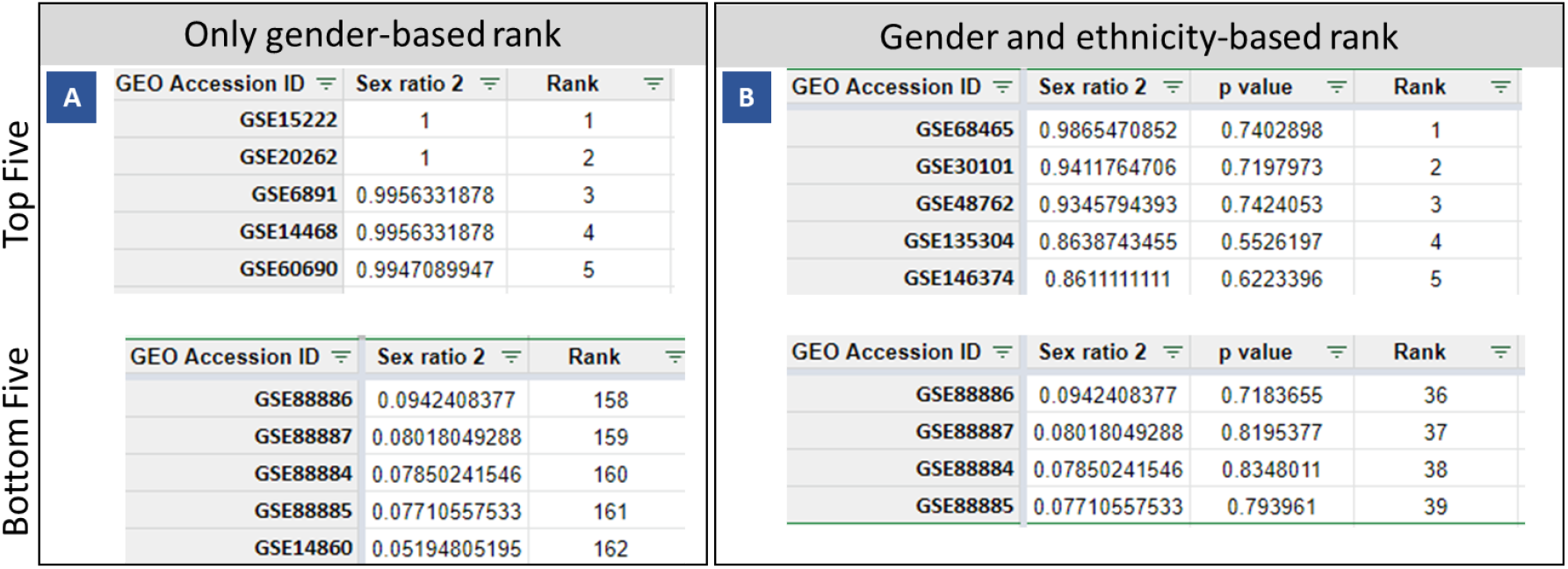
GEO data ranking

#### Ranking GEO datasets based on gender as well as ethnicity/race

To add additional criteria for ethnicity/race, we first extracted the number of individuals labeled for each ethnicity/race mentioned in the metadata. To note, for the purpose of this study, we are considering ethnicity and race the same. To compute an overall score, keeping in mind both gender and ethnicity, we only looked at datasets that contained both information. Then we computed the proportion of each ethnicity for that dataset. This gave us a probability of each ethnicity in the dataset. Next, we employed the Chi-square goodness of fit test to estimate a p-value for each data series. To note, since not every data had all ethnicities/races or the same ones therefore while calculating the chi-square test we made sure we were only looking at the ethnicities mentioned in the dataset under observation and ignored the rest. The expected null hypothesis is that there is no difference between observed and expected probabilities i.e, data does follow a distribution with certain proportions. Each ethnicity should exist in equal proportions in the dataset for unbiased evaluation. Therefore, under these assumptions, the Chi-square goodness of fit test gave us a p-value for each dataset. The larger the p-value, the higher the chance of the null hypothesis being true, and the more balanced, unbiased the data. Next, to compute the overall rank for each dataset, we used the gender ratio (as mentioned above) and the p-value for ethnicity/race. The larger the gender ratio and the higher the p-value the more balanced/unbiased the data. To calculate the rank based on two different criteria, we employed Excel’s rank equation and countif functions. The dataset for the top and bottom 5 datasets can be viewed in **Figure 2B**.

### Evaluating GEO datasets before and after 2015 for DEI criteria

While collecting DEI-relevant information from GEO datasets, we also extracted the corresponding year of data submission from the metadata information. The year information can be employed to compare DEI bias scores for datasets before and after 2015. **Figure 3A-B** shows the changes in the gender ratio and the p-value, however, a closer look at the distribution of the two shows that for some patients after 2015, their gender ratio is more biased compared to their ethnicity ratio. This is indicated in **Figure 4A-B**. Due to the high variability in the scientific question that each study/dataset was addressing it is hard to say anything for sure. However, a closer inspection of a few of the datasets indicates that there are more women recruited for certain studies might be because women are generally higher in number in the world compared to men or because men mostly have financial priority and prefer not to take part in such studies or it can be due to gender disparity in science which is still very common or it can be because of the relatively small size of the datasets under inspection.

**Figure 3.**
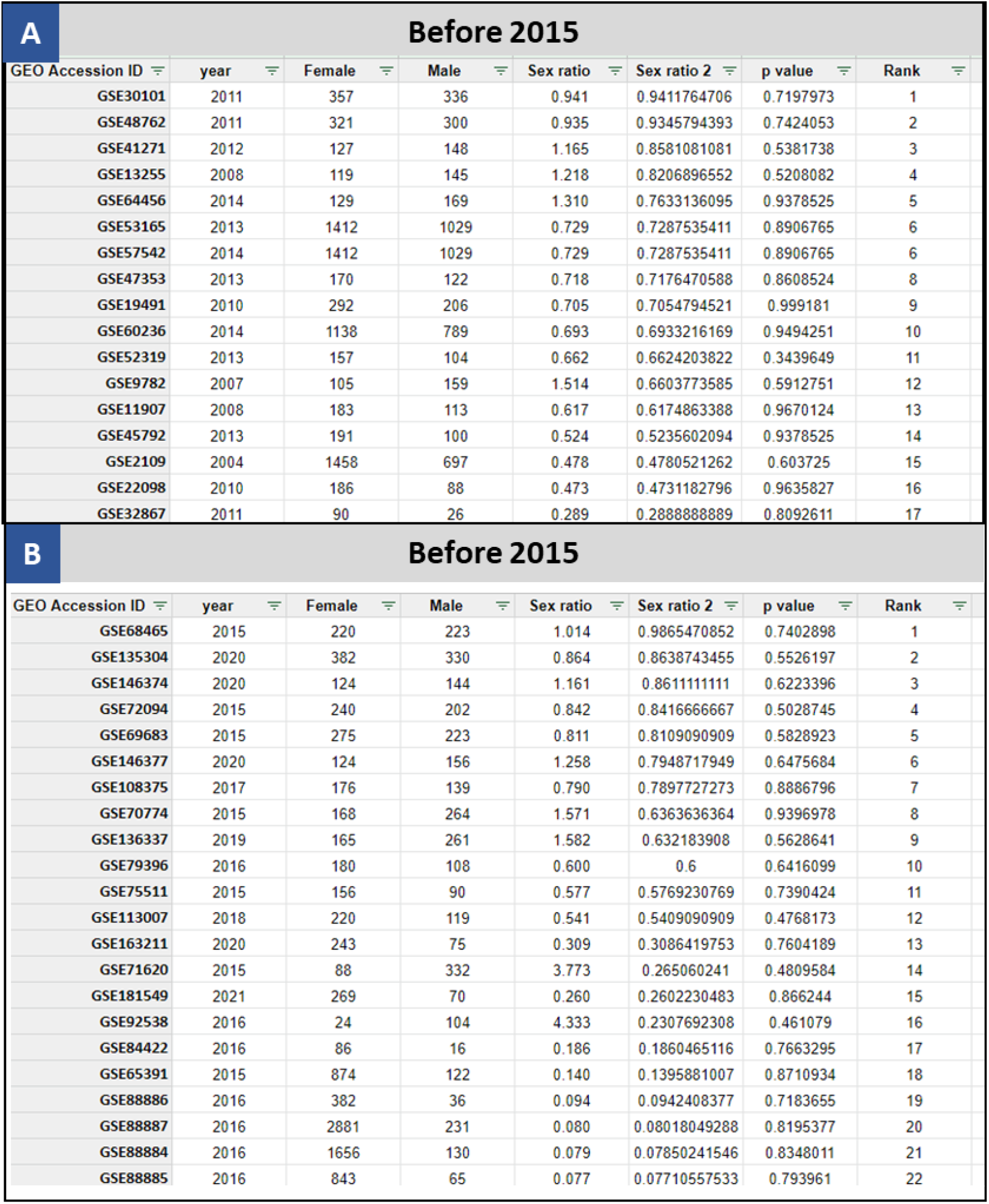
GEO Datasets before and after 2015

**Figure 4.**
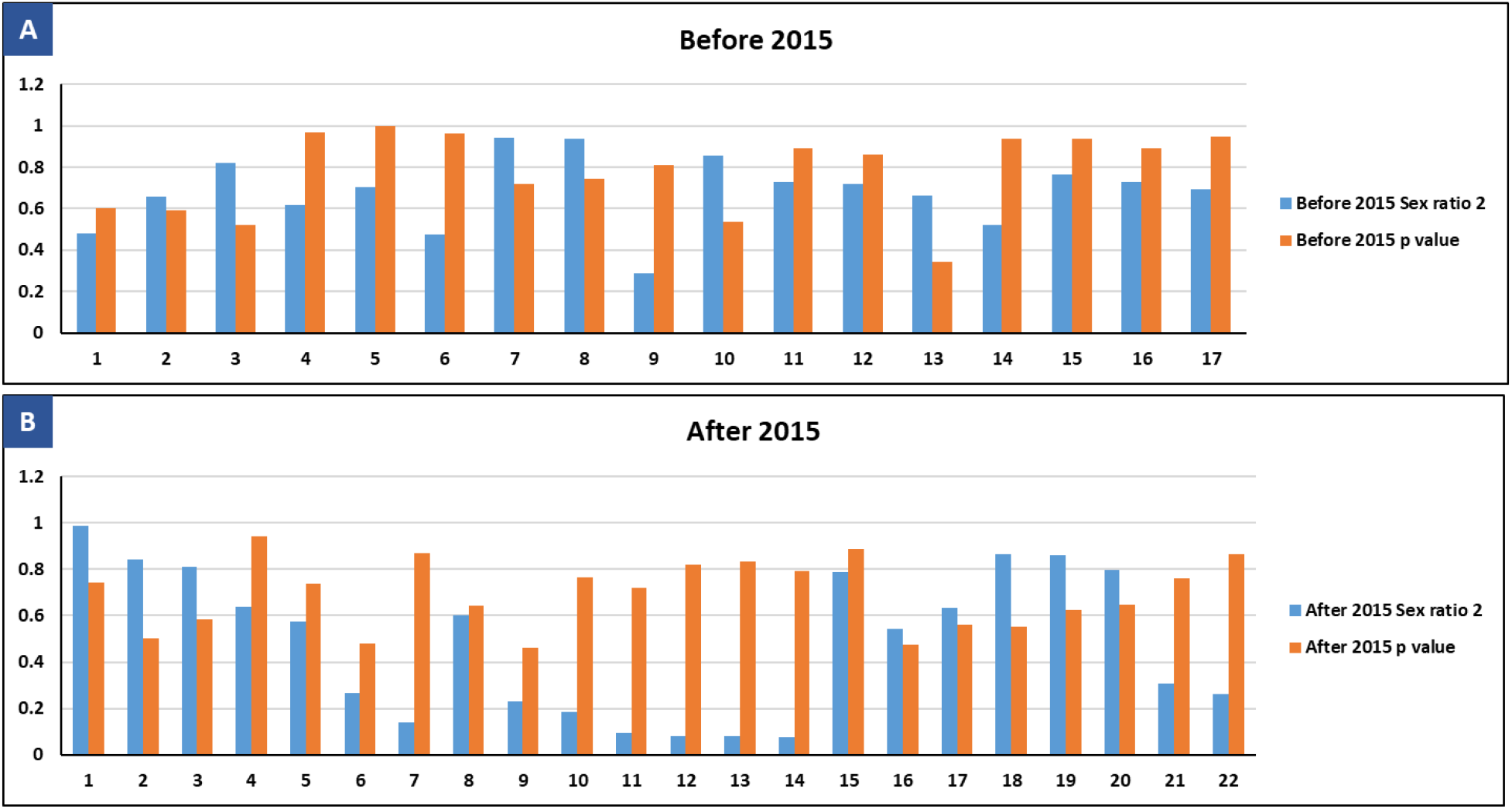
Distribution of gender ratio and p-value for before and after 2015

### 3 Performing exploratory analysis on curated data

A general overview of the curated datasets (**Figure 5**) shows that most of our data is biased with the following ethnicities white, caucasian, African American, and Hispanic.

**Figure 5.**
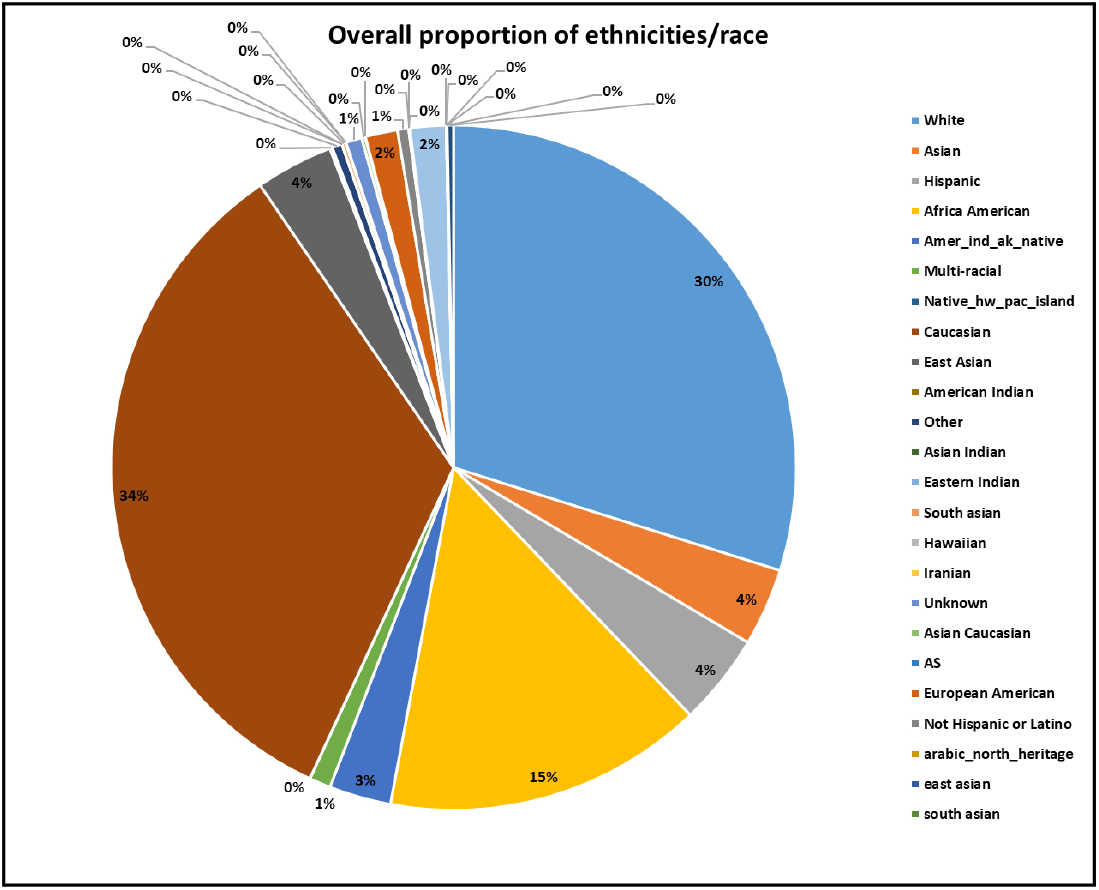
Distribution of ethnicities/race in the curated data

## Conclusion

In this study, we successfully extracted, filtered, and curated a dataset to investigate if Gene Expression Omnibus (GEO) datasets are biased relative to the US population using DEI criteria (gender, ethnicity, ancestry, race). We encountered issues pertaining to how different research groups classify different ethnicities and genders, to make sure we have good data entries, all 211 datasets result was manually examined and curated. Our results “*gender-only ratio”* show that there are 116 datasets from 163 which are equal to and above 0.5 indicating a good balance of gender, whereas 43 were below 0.5 showing gender inequality. From visual inspection, we can see that there are more female samples. This can be because we are only looking at a subset of the entire 583 datasets. Moreover, literature reports do indicate that there has been active recruitment of women in clinical settings ^16^, and the need for gender equality is now more seen than in previous times, although we still have a lot more road to cover. For our “*gender and ethnicity-based overall rank*” we observe that out of our 39 samples that have both gender and ethnicity data, we have a high p-value and gender ratio. This shows that we fail to reject null and our observed and expected ratio of ethnicities are similar and the high gender ratio indicates that our data is more neutral towards gender. However, this is an initial analysis of a limited dataset, and a more broad approval needs to be taken especially while collecting datasets and computing the overall rank. Fisher test can perhaps give better results than the chi-square test, we can also try to include additional DEI criteria in our analysis, and we can also try to investigate those studies that show a very high or very low score the study might itself be biased or they might have overlooked while recruiting their samples. This analysis, on the whole, indicates that there is a need to keep DEI in mind while designing, planning, and executing scientific endeavors, such biases cannot only lead to unjust result outcomes but can also be detrimental in policy and legislature when used in decision-making settings as ground truth. As researchers and scientists we need to develop universal criteria of acceptance, diversity, inclusion, and equality in health science, machine learning techniques, and artificial learning so we do not overlook such important indicators/criteria in our datasets.

## Data availability

The datasets and materials supporting the findings of this study are available in the following repositories. Raw and processed data, as well as formulas used in the analysis, are accessible through the Google Spreadsheet. Additionally, the data, R code, and figure materials can be found in the Google Drive repository.

## References

1. Barrett, T. et al. NCBI GEO: archive for functional genomics data sets--update. Nucleic Acids Res. 41, D991–5 (2013).

2. Gondal, M. N. & Chaudhary, S. U. Navigating Multi-Scale Cancer Systems Biology Towards Model-Driven Clinical Oncology and Its Applications in Personalized Therapeutics. Front. Oncol. 11, 712505 (2021).

3. Khurshid, G. et al. A cyanobacterial photorespiratory bypass model to enhance photosynthesis by rerouting photorespiratory pathway in C3 plants. Sci. Rep. 10, 20879 (2020).

4. Gondal, M. N. et al. TISON: a next-generation multi-scale modeling theatre for in silico systems oncology. Systems Biology (2021).

5. Clough, E. & Barrett, T. The Gene Expression Omnibus database. Methods Mol. Biol. 1418, 93–110 (2016).

6. Patra, B. G. et al. A content-based literature recommendation system for datasets to improve data reusability - A case study on Gene Expression Omnibus (GEO) datasets. J. Biomed. Inform. 104, 103399 (2020).

7. Hadley, D. et al. Precision annotation of digital samples in NCBI’s gene expression omnibus. Sci. Data 4, 170125 (2017).

8. Popejoy, A. B. & Fullerton, S. M. Genomics is failing on diversity. Nature 538, 161–164 (2016).

9. Need, A. C. & Goldstein, D. B. Next generation disparities in human genomics: concerns and remedies. Trends Genet. 25, 489–494 (2009).

10. Gondal, M. N. et al. A personalized therapeutics approach using an in silico Drosophila Patient Model reveals optimal chemo- and targeted therapy combinations for colorectal cancer. Front. Oncol. 11, 692592 (2021).

11. Bentley, A. R., Callier, S. & Rotimi, C. N. Diversity and inclusion in genomic research: why the uneven progress? J. Community Genet. 8, 255–266 (2017).

12. Sirugo, G., Williams, S. M. & Tishkoff, S. A. The missing diversity in human genetic studies. Cell 177, 26–31 (2019).

13. Gondal, M. N., Cieslik, M. & Chinnaiyan, A. M. Integrated cancer cell-specific single-cell RNA-seq datasets of immune checkpoint blockade-treated patients. bioRxivorg (2024) doi:10.1101/2024.01.17.576110.

14. Gondal, M. N., Shah, S. U. R., Chinnaiyan, A. M. & Cieslik, M. A systematic overview of single-cell transcriptomics databases, their use cases, and limitations. Front. Bioinform. 4, 1417428 (2024).

15. Bao, Y. et al. Targeting the lipid kinase PIKfyve upregulates surface expression of MHC class I to augment cancer immunotherapy. Proc. Natl. Acad. Sci. U. S. A. 120, e2314416120 (2023).

16. Liu, K. A. & Mager, N. A. D. Women’s involvement in clinical trials: historical perspective and future implications. Pharm. Pract. (Granada) 14, 708 (2016).

